# Patterned expression of Pannexin 1 channels in the adult cerebellar Purkinje cells

**DOI:** 10.1101/453738

**Authors:** Visou Ady, David Dubayle, Pascale Le Blanc, Valery Shestopalov, Claude Meunier, Carole Levenes

## Abstract

Pannexin1 (PanX1) are recently discovered proteins that can form large pore channels at the cell surface. They have been implicated in ATP-dependent cell-to-cell communication and in several pathophysiological processes such as inflammation, cell death and epilepsy. Using immunohistochemistry in the adult mouse, we describe the presence of PanX1 in the deep cerebellar nuclei, in large cells of the granular layer, presumably Golgi interneurones, in some Bergmann glia radial processes as well as in the soma and dendrites of Purkinje cells. In the latter, PanX1, like many other proteins, distribute heterogeneously. Only Zebrin II-positive Purkinje cells express PanX1, in accordance with the so-called “zebra-striped” modular architecture of the cerebellum. This distribution in zebra-stripes suggest that PanX1 may contribute to the control of ensemble activity within cerebellar microdomains or to the response of Purkinje cell to excitotoxicity and cell-death messages.

## Introduction

Pannexins are membrane proteins ubiquitously present in vertebrate organisms. They were discovered in 2000 by sequence homology with the invertebrate innexins [1]. They show structural similarities with connexins [1] but do not seem to form gap-junctions in native cells ([2] and refs in [3]. Among the three types of pannexins described to date, pannexin 1 (PanX1) is by far the most studied. It is heavily expressed in the central nervous system, in both neurons and glial cells. Its mRNA has been detected in the retina, the cerebellum, the neocortex and hippocampus, in the amygdala, *substantia nigra* and the olfactory bulb, [4-6, 7, 8, 9, 10, 11]. PanX1 can be activated by cytoplasmic Ca^2 +^, membrane depolarization, extracellular ATP and K^+^, by mechanical stretch and caspase cleavage ([12, 13, 14] and refs in [3]). They can form large conductance channels of up to 550 pS [13] that carry non-selective ion fluxes and are permeable to molecules up to 1 kDa, including ATP itself [14, 15],[16]. Once released, ATP can act on purinergic receptors or can be further degraded into adenosine by ectonucleotidases. Thus pannexins mediate auto- and paracrine communication, making them possible actors of retrograde neuronal communication and modulators of neuronal activity, for example through presynaptic adenosine receptors [17]. However, their exact physiological role remains unclear. Evidence shows that they can initiate Ca^2+^ waves, for instance after mechanical stretch [13], control the vascular tonus, regulate differentiation, cell death and immune function [3]. In pathological conditions, they may activate the inflammasome ([12] reviewed in [18]), contribute to caspase-mediated apoptosis [19] and ischemic cell-death [20], and enhance epileptic seizures. They could also play a role as tumor suppressors (refs in [3]).

In the cerebellum, PanX1 proteins are heavily expressed, particularly in Purkinje cells [7, 10], but their contribution to cerebellar physiology or pathology remains totally unknown. Despite its apparent homogeneous crystal-like organization, the cerebellum is, in fact, highly compartmentalized. It displays genetically determined and reproducible topographic domains: the transverse zones and parasagittal stripes, including the cortical so-called zebra-stripes [21]. The latter can be revealed by the patterned expression of numerous molecules, particularly in Purkinje cells. Zebrin II, now known to be the respiratory enzyme Aldolase C, was the first example of such a protein [22]. It is expressed in parasagittal bands of Purkinje cells that extend throughout the cerebellar cortex, separated by bands of Zebrin II-negative Purkinje cells. Many other proteins match Zebrin II Purkinje cells patterns. For example, phospholipase C β4 (PLC β4; [23]), the metabotropic glutamate receptor 1b (mGluR1b; [24]) and the early B-cell factor 2 (EbF2; [25]) are expressed in Zebrin II-negative Purkinje cell stripes. On the other hand, PLCβ3 [26], the excitatory amino-acid transporter 4 (EAAT4; [27]), and the GABA-B 1b receptor [28] are expressed in Zebrin II-positive Purkinje cell. These striped distributions of neurotransmitter receptors and their effectors indicate that Zebrin II-negative and -positive Purkinje cells respond differentially to their inputs. For instance, the glutamatergic climbing fibers that form synapses onto Zebrin II-positive Purkinje cells release more glutamate per action potential than those terminating on Zebrin II-negative ones [29]. Zebrin II-positive Purkinje cells seem to be adapted to respond to increased glutamate inputs and to better resist death and neurodegeneration of various origins [30, 31, 32, 33]. Sphingosine kinase, a lipid metabolism protein that plays an important role in apoptosis, is also found selectively in Zebrin II-positive Purkinje cells [30]. Overall, the proteins expressed in Zebrin II-positive Purkinje cells, including Zebrin II itself, seem to confer these cells capacities to respond to sustained neuronal activity and to the associated increase in metabolic demand.

In the present study, we use a type of antibodies directed against the extreme C-terminal domain of PanX1 to investigate the distribution of PanX1 in the cerebellum. These antibodies that had never been used in this structure reveal a zebra-striped pattern of PanX1 in Purkinje cells. This differential rostro-caudal expression,typical of the cerebellum, had never been described before for PanX1. It suggests that PanX1s contribute to the adaptation of Purkinje cells to the increased activity observed in the zebra stripes, and thus that pannexins may play important functions in the physiology and pathophysiology of the cerebellum.

## Materials and methods

### Animals

The animals used were all adult (n=25) C57BL/6JRj males and females C57BL/6JRj mice (from 8 to 12 weeks of age), purchased at Janvier Laboratory (Le Genest-St-Isle, France). Mice were housed at the central animal facility of Les Saints-Pères (Paris, France). Housing and all procedures were performed in accordance with the guidelines of the French Ministry of Agriculture and the European Community and have been approved by the ethical committee of Paris Descartes University (agreement: Paris Descartes # 14-013). A minimal number of animals was used and they were handled with maximum care in order to minimize their suffering.

### Tissue preparation

The animals were deeply anaesthetized with sodium pentobarbital (175 mg/Kg i.p.). The intracardiac perfusion consisted of 50 ml 0.01 M phosphate-buffered saline (PBS, pH: 7,4), followed by 50 ml of 4% paraformaldehyde in 0.1 M PBS. The cerebellum was removed, postfixed for 1 day in 4% paraformaldehyde solution and cryoprotected in a 30% sucrose phosphate-buffered solution at 4°C for 2 days before cutting. The cerebellum was cut into longitudinal or sagittal 80 mm-thick sections using a freezing sliding microtome (Frigomobil Reichert-Jung) and collected in plastic-wells containing PBS. Serial sections over the entire rostrocaudal length of the cerebellum were treated and analyzed.

### Antibody Characterization

**Table 1.** Primary antibodies used in this study.

| Antigen | Immunogen | Source, host species, RRID | Dilution |
| --- | --- | --- | --- |
| **Pannexin 1** | 18 kDa C-terminal portion of human Pannexin 1 protein | Chemicon, Millipore, USA ; rabbit polyclonal ; affinity purified.<br>RRID : AB_2236635 | 1:750 |
| **Pannexin 1** | 18 kDa C-terminal portion of human Pannexin 1 protein, affigel-conjugated | Dr. V. Schestopalov, Miami, USA ; rabbit polyclonal ; affinity purified.<br>RRID : AB_2315054 | 1:750 |
| **Calbindin** | Calbindin D-28k protein | Swant, Switzerland ; mouse monoclonal.<br>RRID : AB_10000347 | 1:7500 |
| **Zebrin II (Aldolase C)** | Recognizes a single polypeptide band of apparent molecular weight 36 kDa on cerebellum homogenates from all vertebrate classes studied with this Ab. | Dr. R. Hawkes, Calgary, Canada ; mouse monoclonal.<br>RRID : AB_2315622 | 1:50 |

The two types of PanX1 antibodies that we used were rabbit polyclonal antibody raised against the C-terminus of the human PanX1. The first one came from Chemicon (USA). The second one was made by V. Schestopalov [11] as follows. A rabbit polyclonal antibody against the carboxyl terminus of human PanX1 cDNA encompassing the entire coding region was synthesized by PCR amplification of the cDNA insert from clone IMAGE: 4390851 using two gene-specific primers (Px1F: 5′-TCTGGATCCTACACGCTGTTTGTTCCA-3′; Px1R: 5′-TCTAAGCTTGCAAGAAGAATCCAGAAG-3′). It was then inserted into the BamHI and HindIII sites of pET-23a to yield the pETPx1 plasmid. The resultant 18 kDa pETPx1 protein fused to the His-tagged C-terminus was purified from E. coli and used for immunization of the rabbit. The affinity purified rabbit serum containing specific anti-human PanX1 activity and lacking any significant non-specific activities was used for the immunohistochemistry. In the present study, we verified the specificity of the two PanX1 antibodies we used with their immunogenic peptide in western-blot and immunohistochemistry experiments (see results). The sequence of the peptide corresponds to the 135 last amino-acids (C-ter) of the human PanX1 protein. Blasting this peptide sequence shows its high specificity for PanX1. The specificity of PanX1 antibody was tested by peptide pre-adsorption: 1/2000 anti-PanX1 was pre-incubated for 2 h at room temperature with 1/400 of immunizing peptide and then used for western-blot of 30 µg of protein extract from adult mice cerebellum. The same peptide was used in immunohistochemistry with the same pre-incubating procedure except that the PanX1 antibody was used at 1/750 and blocking peptide dilutions were 1/750, 1/150 and 1/75. Both anti-PanX1 antibodies gave similar results. Therefore, data obtained with these to different antibodies were pooled.

### Immunostaining for light microscopic analysis

Cerebellar sections were processed for PanX1, calbindin (CaBP), Zebrin II expression analysis. Sections were first incubated in blocking buffer containing PBS-T-G (PBS 0.1%, Triton X-100 0.25%, Glycine 0.75% and gelatine from porcine skin 0.25%, Sigma, UK) for 1 hour at room temperature. Following three washes in PBS-G (PBS-T-G, without Triton), sections were incubated for 72 hours at room temperature in a mixture of the primary antibodies. Following three washes in PBS-G, the slices were incubated at room temperature for 2 hours with a mixture of secondary antibodies containing: anti-mouse Alexa fluor (AF) 546 (1:250) and anti-rabbit AF488 (1:250), both from Molecular Probes (USA). Following a final wash in PBS-G, the sections were air-dried and mounted in a homemade Mowiol-based medium. Images were acquired with a LSM510 or LSM710 microscope (Zeiss) and processed with the ImageJ software (public domain: http://rsbweb.nih.gov/ij/). Final montages were made with Adobe Photoshop.

## Results

### Distribution of PanX1 immunoreactivity: rostrocaudal gradient and parasagittal stripes

We first verified the specificity of the PanX1 antibodies using western blot and immunohistochemistry. In western blots of mouse cerebellar extracts, the PanX1 antibodies recognize a single ~ 43 kDa band as expected for PanX1 in brain extracts [11]. This band disappeared after incubating the antibodies with a saturating concentration of their immunogenic peptide (Fig 1A). In the immunohistochemistry experiments, the pattern of PanX1 labeling observed in sagittal cerebellar sections vanished with increasing concentrations of the blocking peptide in a dose-dependent manner (Fig 1B-E). We also verified that the blocking peptide did not interfere with the immunolabeling procedure by testing it with an antibody directed against parvalbulmin (Fig 1F). As expected, the peptide did not affect the parvalbumin labeling.

**Fig 1.**
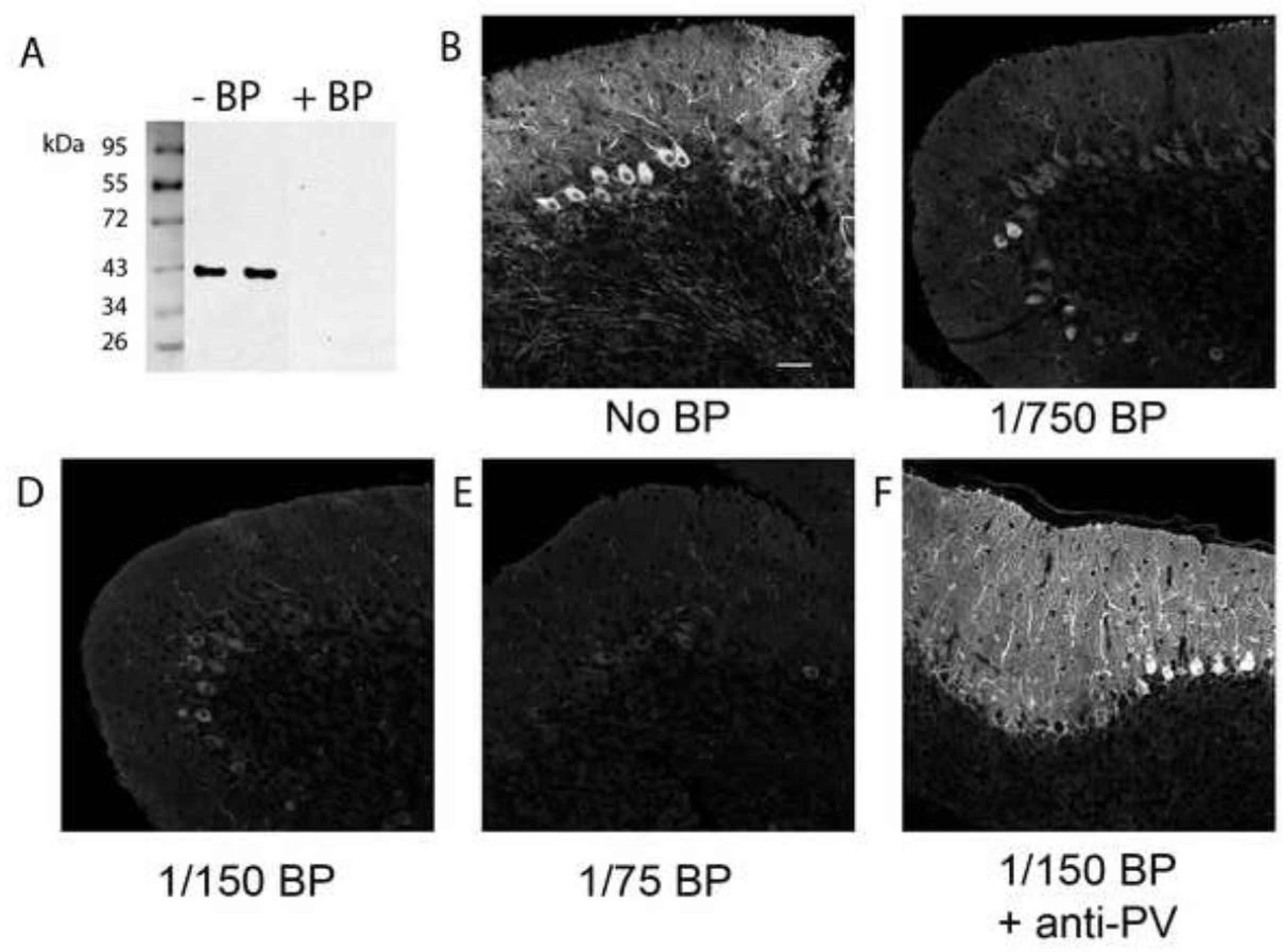
Specificity of the anti-PanX1 antibody.

Western blot of mouse cerebellum extracts labeled with the PanX1-antibody. **A**: The two left lanes correspond to PanX1 antibody alone (- BP); the two right lanes correspond to PanX1 antibody + blocking peptide (+ BP). **B**: Confocal images of immunohistochemical labelings of PanX1 without blocking peptide. **C**-**E**: Confocal images with the same antibody incubated in the presence of increasing concentrations of the blocking peptide (dilutions as indicated). **Images** B to E were taken from Lobule X. **F**: Immunolabeling of parvalbumin (PV) protein made in the presence of the PanX1-blocking peptide (1/150). This control indicates that the blocking peptide does not inhibit any immunohistochemistry reaction but that directed against PanX1. Scale bar: 50 µm.

We next used the anti-PanX1 antibodies to assess the distribution of PanX1 in the cerebellum. As a marker of Purkinje cells, we used the calcium binding protein calbindin-D28k (CaBP). CaBP labeling also allowed us to verify the aspect of these cells and the quality of intracardial fixation. Confocal images of sagittal cerebellar slices made in the vermis revealed that PanX1 antibodies strongly label the soma and dendrites of Purkinje cells (Fig 2A), as previously observed [10, 34]. The labeling was spotty and more concentrated in proximal dendrites (Fig 2A,B). In distal branches, it was also detectable in the vicinity of spines at the level of the thin spiny branchlets (Fig 2B top insets). Remarkably, numerous PanX1 spots seemed to be located outside Purkinje cells themselves, in close apposition to their dendrites. This is suggestive of a possible PanX1-mediated communication between Purkinje cells spines and their afferences (Fig 2B top insets). Molecular layer interneurons were negative for PanX1 labeling (Fig 2B). The radial processes of some - but not all - Bergman glial cells were immunoreactive for PanX1 (Fig 2B). Granule cells were devoid of labeling, but longitudinal sections revealed the presence of PanX1 in round cell bodies of the granular layer, presumably from Golgi cells (Fig 2C). In the deep cerebellar nuclei (DCN), Purkinje cell axons were strongly immunoreactive for PanX1, and some large cells immunopositive for CaBP were also reactive for PanX1 antibodies (Fig 2D). All these observations are in agreement with previous literature [7, 10, 35, 36].

**Fig 2.**
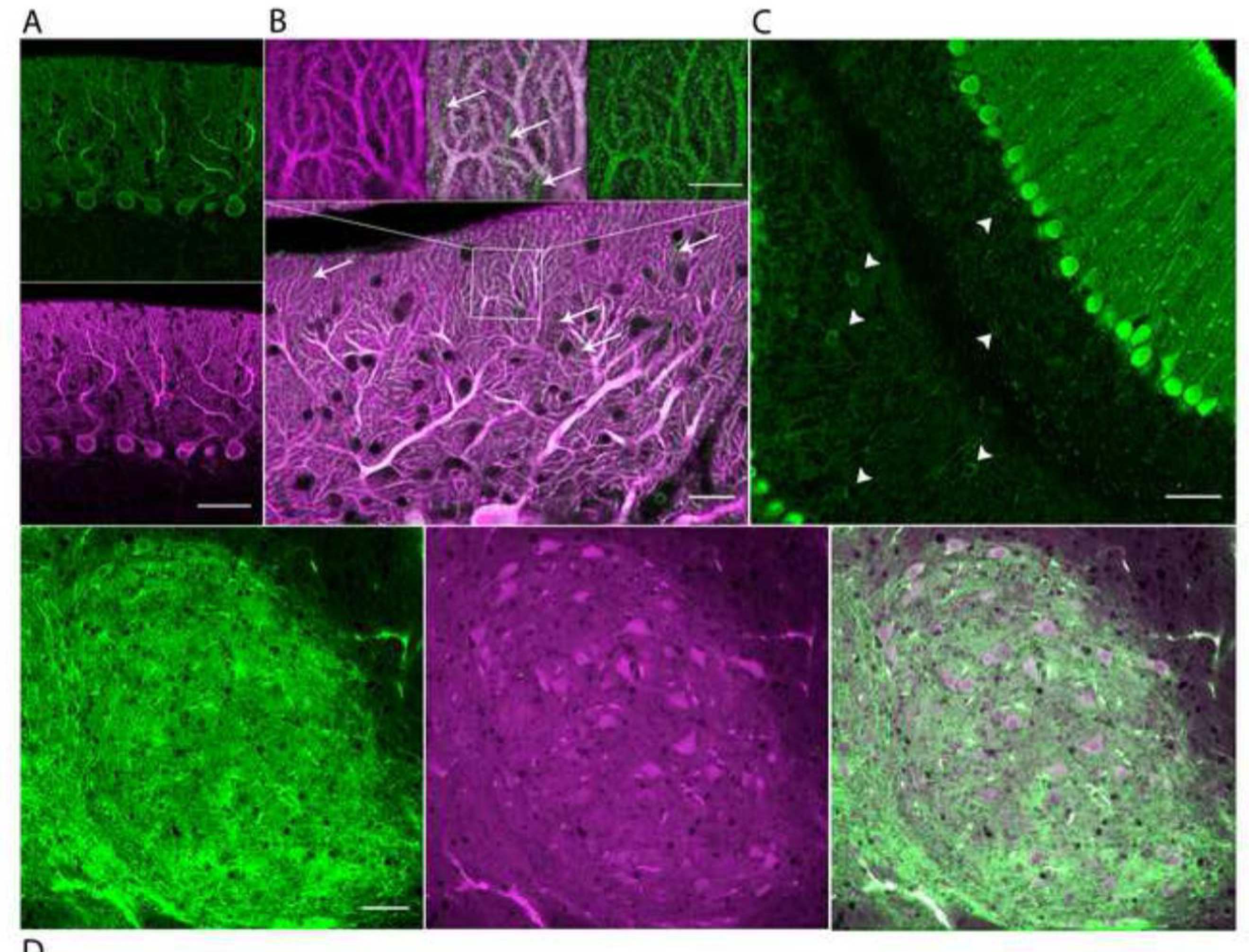
Cellular distribution of PanX1 in the cerebellar cortex.

**A**: Confocal images acquired in a PanX1 positive zone, PanX1 in green (top), CaBP in magenta (bottom). **B**: At high magnification, Purkinje cell dendrites show dense punctiform PanX1 staining, notably in the spiny branchlets (top inserts). Notice the numerous PanX1 positive radial processes that are likely Bergman glial processes (white arrows in merged image, top center and bottom). **C**: Labeling in longitudinal cerebellar sections confirms Purkinje cell staining and reveals round middle size cells immunopositive for PanX1 (white arrows heads), presumably Golgi cells. **D**: Fastigial deep cerebellar nucleus. Some large cells are immunopositive for PanX1 (in green) and for CaBP in magenta (center). Smaller cells are also positive for PanX1 but negative for CaBP (merged image, right). Notice the intense labeling of Purkinje cells axons. Scale bars in A : 50 µm; in B : 20 µm and 10 µm (inserts); C and : 50 µm.

Intriguingly, and in contrast with previous data, PanX1 labeling of Purkinje cells was almost absent in the anterior cerebellar lobules, although well-shaped Purkinje cells were present, as attested by CaBP labeling (Fig 3A). Mosaic parasagittal images of entire cerebellar sections (Fig 3B) revealed a clear rostro-caudal gradient of PanX1 labeling, in contrast to the homogeneous CaBP labeling. This was particularly obvious in the vermis, when comparing the rostral lobule III (Fig 3A) with the caudal lobule IX (Fig 3C). In the central and posterior lobules, PanX1 immunofluorescence was discontinuous, as PanX1-immunopositive zones were intermingled with regions of PanX1-immunonegative Purkinje cells. Staining in the lobule × was constant among slices and more pronounced than in the other lobules.

**Fig 3.**
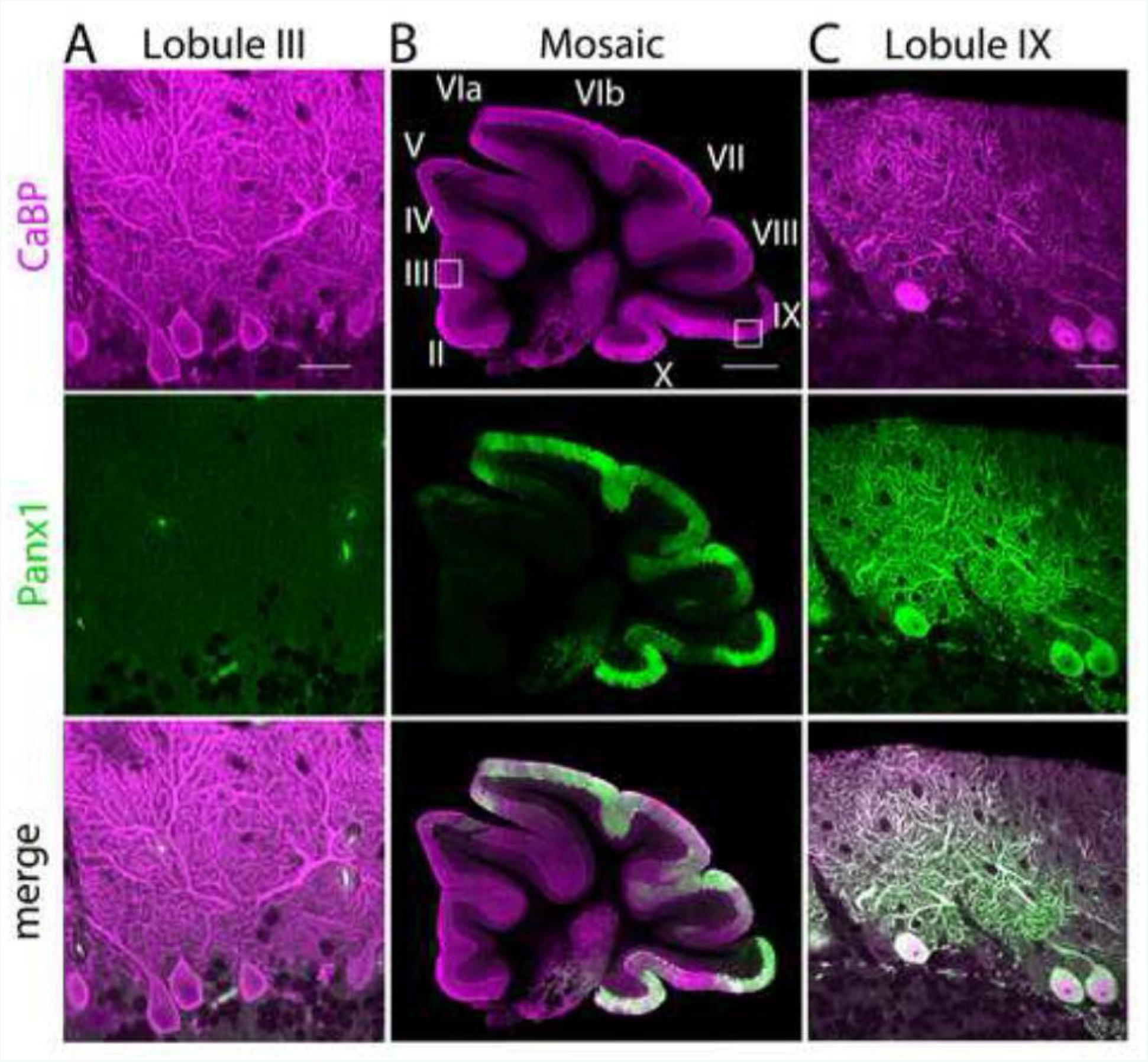
Localization of PanX1 in the cerebellum: dense labeling in Purkinje cells of the posterior lobules.

**A**: In lobule III, molecular layer displays very low PanX1 staining (green), although Purkinje cells are well present (CaBP, in magenta, was used as a Purkinje cell marker). **B:** Automated mosaic confocal images of sagittal slices from the cerebellar vermis region. White squares indicate the location of the high magnification confocal images shown in A (lobule III) and C (lobule IX). **C**: Purkinje cells soma and dendrites in lobule IX display very strong PanX1 labeling. Scale bars in A and B: 20 µm B; in C: 500 µm.

### Distribution of Pannexin1 labeling matches that of Zebrin II

The rostro-caudal heterogeneous expression pattern of PanX1 that we observed in sagittal sections is reminiscent of that of the several tens of proteins expressed in zebra stripes in the cerebellum (reviewed in [21]), the most studied being Zebrin II [22], identified as the Aldolase C enzyme [37]. The very name “Zebrin” comes from the zebra-like pattern that clearly appears in longitudinal sections.,

Labelings of PanX1 in longitudinal sections displayed a striking zebra-type distribution, while that of CaBP appears homogeneous attesting the actual presence of Purkinje cells in the overall slice (Fig 4). In those longitudinal sections, the floculi, parafloculi and Crus II regions consistently appeared PanX1 positive (Fig 4A,D), while lobules I to V displayed several little stripes of few immunopositive Purkinje cells (Fig 4B,C).

**Fig 4.**
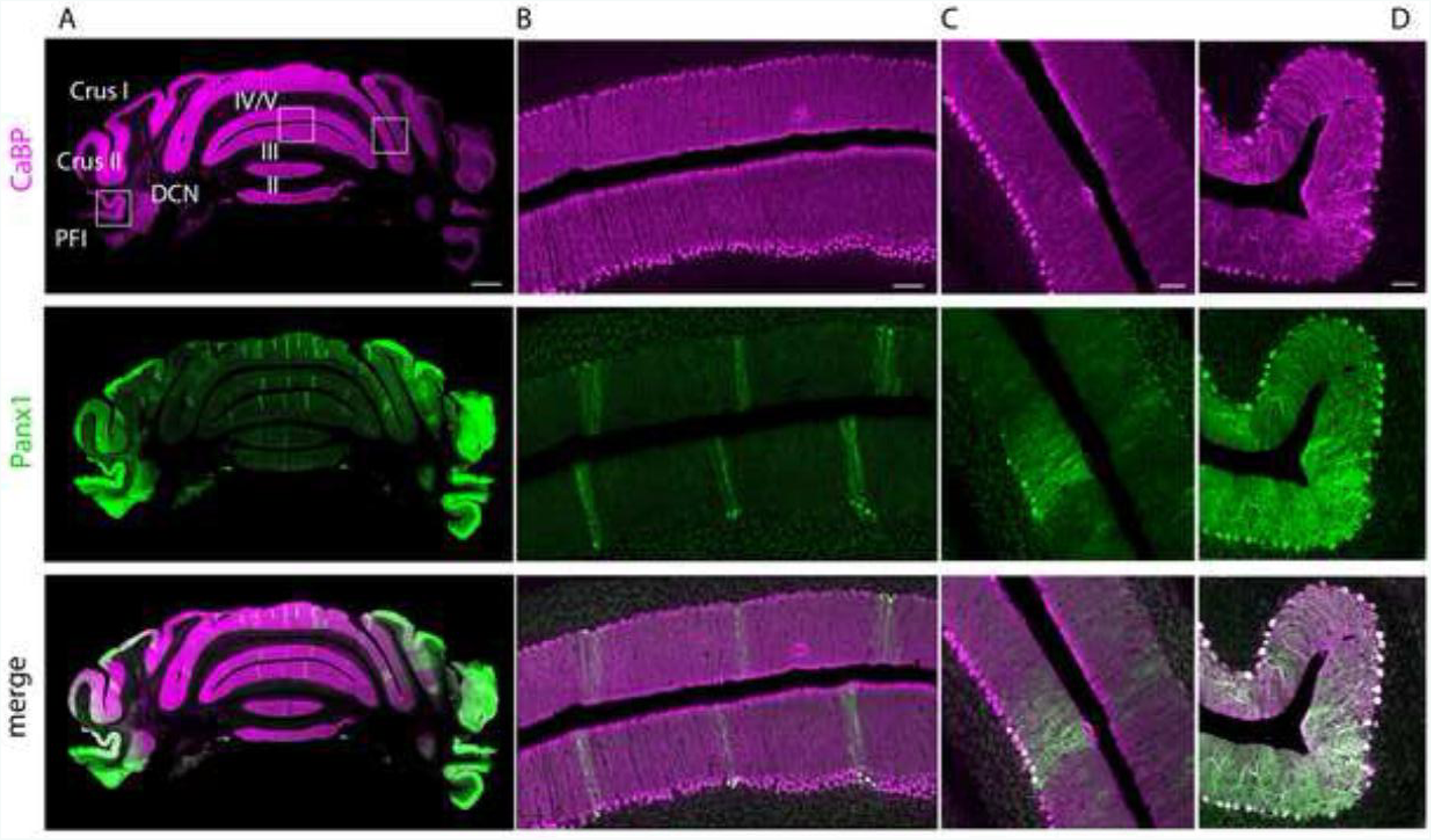
Longitudinal cerebellar sections reveal striped PanX1 labeling.

**A**: Mosaic reconstructions of CaBP (magenta) and PanX1 (green) labeling show that PanX1 are distributed in subsets of several tens of Purkinje cells drawing stripes in lobules I-V. **B**, **C** and **D** are confocal images made in the regions indicated by white squares in A. Note the intense PanX1 labeling in Crus I/II and in the paraflocculi (PFl). DCN, deep cerebellar nucleus, Roman numbers indicate to corresponding lobules. Scale bars in A: 500 µm, B: 100 µm, C and D: 50 µm

The profile of PanX1 distribution in rostral longitudinal section being typical of that of Zebrin II, we made double labelings of these two proteins to compare their respective distributions using large scale mosaic imaging (Fig 5).

**Fig 5:**
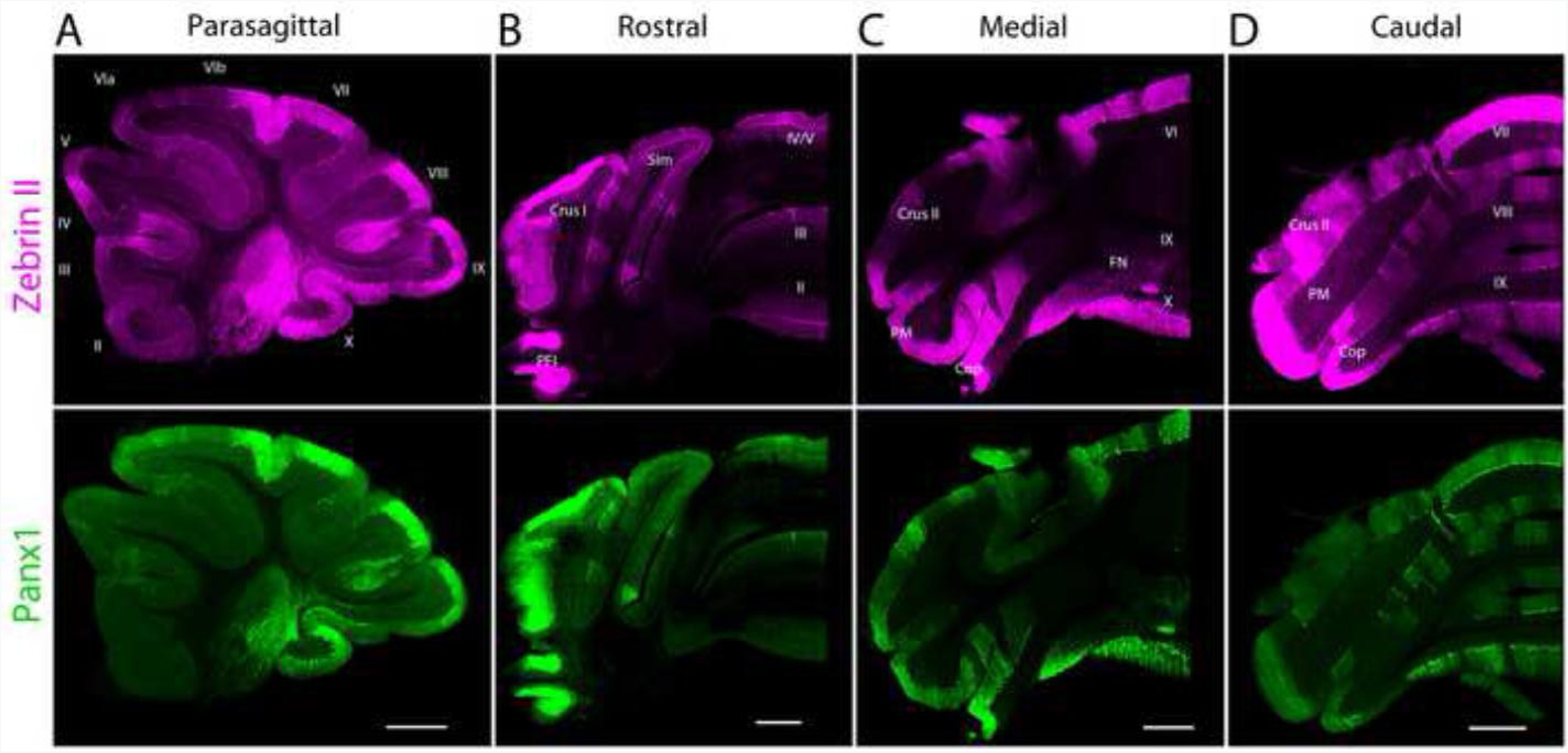
PanX1 and Zebrin II distribution in Purkinje cells match throughout the cerebellum.

Images are mosaic reconstruction made from confocal images. **A**: Zebrin II and PanX1 immunolabeling colocalize in Purkinje cells in a sagittal cerebellar slice taken in the vermis. **B**: Same colocalization observed in a longitudinal slice made in the rostral part of the cerebellum. **C**, **D**: Same as in B but in the medial and caudal parts of the cerebellum respectively. Lobules are indicated as Roman numbers; PFl, paraflocculus; Sim, *lobulus simplex*; Cop, *copula pyramidis*; PM, *lobulus paramedianum*; FN, fastigial nucleus. Scale bars in B to F: 500 µm.

In parasagittal sections, the overall distribution of PanX1 matched that of Zebrin II (Fig 5A). This was clear from lobule VIb to the posterior tip of lobule VII, and in lobules VIII to X, where Purkinje cells where almost uniformly labeled. In other lobules, there was sometimes a difference in the intensity of staining, which was likely due to different subcellular distributions of Zebrin II and PanX1. However, examination at higher magnification showed that the boundaries of strongly immunoreactive bands and weakly immunoreactive (or unreactive) bands were very similar for these two markers.

We made longitudinal sections in the rostral, medial and caudal parts of the cerebellum to verify if PanX1 and Zebrin II distributions matched over the whole antero-posterior structure (Fig 5B-D). In rostral regions, most Purkinje cells were Zebrin II-negative and PanX1-negative (Fig 5B). In the vermis, the midline and the two lateral Zebrin II-positive bands on each side (presumably P1+, P2 +, P3+ according to [38]) were PanX1-positive and were separated by broad PanX1-negative bands (lobule III or IV). In medial and caudal sections, staining in lobules VIb, VII and × was uniform and dense across Purkinje cells (Fig 5C and D). In lobule IX, the midline and the two lateral Zebrin II-positive bands were also PanX1-positive (Fig 5D). Sections through the posterior and the nodular cerebellum showed perfect match between PanX1- and Zebrin II-positive Purkinje cells (Fig 5D).

In conclusion, the distribution of PanX1-positive Purkinje cells matches that of Zebrin II-positive cells either in rostral, medial or caudal parts of the cerebellar cortex.

## Discussion

In the present study, we detect PanX1 in Purkinje cells, DCN cells, Bergmann glia processes and in scattered cells in the granular layer (likely Golgi interneurons) [7, 9, 10, 35, 36]. Our results are in agreement with previous literature for all cell types expressing PanX1. However, we evidence for the first time a rostro-caudal gradient in the distribution of PanX1 in cerebellar Purkinje cells. In our experiments, PanX1 were almost undetectable in the anterior lobules but strongly labeled in the caudal lobules, particularly in the posterior zone. This distribution matches that of the standard in the subject, the Zebrin II protein, which is expressed according to a striped pattern, highly reproducible and highly conserved across evolution [21]. To our knowledge, the antibodies that we used here had only been tested before in the retina [11], but never in the cerebellum. In our western-blots, made on adult cerebellum protein extracts, the anti-PanX1 antibodies detect a 43 kDa band as expected for PanX1. In addition, we used the corresponding immunizing PanX1 peptide pre-adsorbed and incubated with the PanX1 antibody to verify specificity, both in western-blot and in immunohistochemistry. In this latter set of experiments, the dose-dependent blocking effect of the immunizing peptide further confirmed the validity of our data.

Why the specific localization of PanX1 in Zebrin II-positive Purkinje cells has never been reported before is an intriguing question. It may be that our antibodies recognize a different subset of PanX1 isoform(s) compared to previous studies. Several protein modifications explain the various banding patterns observed in western-blots, such as pre- and post-translational modifications (glycosylation, S-nitrosylation or caspase-mediated cleavage). PanX1 seems to be subjected to numerous splicing modifications. Besides the full-length variant named PanX1a (48 kDa), alternative splicing of the human or rat PanX1 encoding genes generates at least four variants (PanX1a-d), of predicted molecular weights of 48, 40, and 34 kDa for PanX1a, 1c, and 1d respectively [39]. PanX1 can also be glycosylated and S-nitrosylated, which modulates their surface expression and activity [34, 40]. This could explain discrepancies between our study and the others published so far. Cone *et al.* [34] made an elegant comparative analysis of labelings obtained with four different antibodies in the adult mouse cerebellum, in which they show that different polyclonal antibodies actually detect distinct species of PanX1. These antibodies revealed various sub-cellular and cellular localization of PanX1, depending on the epitope design. Our antibodies likely target a specific species that is restricted to Zebrin II-positive Purkinje cells. PanX1 isoforms with molecular weight around 43 kDa, like ours, likely corresponds to non - or weakly - glycosylated forms that tend to be more confined to intracellular compartments [34] but can also reach the membrane in some conditions [34, 41]. The species that we detect here seems to reach the membrane because dendrites and spines were heavily stained in our immunohistochemistry experiments.

The distribution of PanX1 proteins in zebra stripes is intriguing,. PanX1 have been so far essentially implicated in pathologies. They have been shown to be activated by pathological conditions such as epileptic seizures or infection [42, 43]. In turn, once activated, they contribute to worsen these conditions. Their activation can engage the inflammasome and trigger aptototic processes [44, 45] (see also [46]). Zebrin II-positive Purkinje cells better resist to infectious and chemical injuries than their Zebrin II-negative sisters and resist better to cell-death. Therefore, the preferential localization of PanX1 in Zebrin II bands seems rather paradoxical, unless the specific species of PanX1 we have evidenced there display a higher threshold for activation. In this case, they would be less activated in pathological condition than their Zebrin-II negative counterparts, thus rendering the Zebrin-II positive cells more resistant to insults. However, this remains an unsubstantiated hypothesis.

Another picture emerges if we consider instead the role of pannexins in normal conditions [3], rather than focus on their deleterious effects. PanX1 exert some of their paracrine functions as ATP channels. ATP, which in turn activates PanX1, can be hydrolyzed into ADP, AMP, and eventually in adenosine by ectonucleotidases. These molecules are agonists of purinergic and adenosine receptors, known to modulate neuronal activity in the cerebellar cortex [47, 48]. For example, adenosine receptors A1R, expressed at both pre- and post-synaptic sides glutamatergic afferences on Purkinje cells, decrease glutamate release [49] and inhibit long-term depression [50]. Thus pannexins could trigger the activation of presynaptic A1R by releasing ATP - further converted into adenosine - and reduce glutamate release. In this scheme, PanX1 would participate to a negative feedback control of synaptic transmission. Such a mechanism would provide a homeostatic regulation of neuronal transmission in the cerebellum, as has recently been suggested elsewhere [51]. The presence of PanX1 in Zebrin II-positive regions could thus help neurons adapt to the increased activity observed in these microzones. It is striking that the majority of the molecules present in parasagittal bands are directly involved in synaptic transmission, control of neuronal activity, or cell responses to the associated metabolic demand. The specific species of PanX1 present in zebrin II-bands may well contribute to this adaptation.

In conclusion, we can hypothesize that the PanX1 species we have evidence here provide Zebrin II-positive Purkinje cells with additional tools to adapt to the specific physiology of this zone characterized by increased glutamate release, enhanced metabolic demand and better resistance to cell-death. All these possibilities remain hypotheses for the moment, but our data open new perspectives that call for further investigation.

## Acknowledgements

We thank Richard Hawkes for the kind gift of the anti-Zebrin II antibody as well as precious recommendations. We also thank David Hansel for critical reading and comments on the manuscript. We sincerely acknowledge the imaging facility SCM (Service Commun de Microscopie - Faculté des Sciences Fondamentales et Biomédicales - Paris) and specially Jean-Maurice Petit.

